# The seasonality of diarrheal pathogens: A retrospective study of seven sites over three years

**DOI:** 10.1101/541581

**Authors:** Dennis L. Chao, Anna Roose, Min Roh, Karen L. Kotloff, Joshua L. Proctor

## Abstract

**Background:** Pediatric diarrhea can be caused by a wide variety of pathogens, from bacteria to viruses to protozoa. Pathogen prevalence is often described as seasonal, peaking annually and associated with specific weather conditions. Although many studies have described the seasonality of diarrheal disease, these studies have occurred predominantly in temperate regions. In tropical and resource-constrained settings, where nearly all diarrhea-associated mortality occurs, the seasonality of many diarrheal pathogens has not been well characterized. As a retrospective study, we analyze the seasonal prevalence of diarrheal pathogens among children with moderate-to-severe diarrhea (MSD) over three years from the seven sites of the Global Enteric Multicenter Study (GEMS), a case–control study. Using data from this expansive study on diarrheal disease, we characterize the seasonality of different pathogens, their association with site-specific weather patterns, and consistency across study sites.

**Methodology/Principal Findings:** Using traditional methodologies from signal processing, we found that certain pathogens peaked at the same time every year, but not at all sites. We also found associations between pathogen prevalence and weather or “seasons”, which are defined by applying modern machine-learning methodologies to site-specific weather data. In general, rotavirus was most prevalent during the drier “winter” months and out of phase with bacterial pathogens, which peaked during hotter and rainier times of year corresponding to “monsoon”, “rainy”, or “summer” seasons.

**Conclusions/Significance:** Identifying the seasonally-dependent prevalence for diarrheal pathogens helps characterize the local epidemiology and inform the clinical diagnosis of symptomatic children. Our multi-site, multi-continent study indicates a complex epidemiology of pathogens that does not reveal an easy generalization that is consistent across all sites. Instead, our study indicates the necessity of local data to characterizing the epidemiology of diarrheal disease. Recognition of the local associations between weather conditions and pathogen prevalence suggests transmission pathways and could inform control strategies in these settings.

## Introduction

Pediatric diarrheal disease is caused by a wide variety of pathogens [1–3]. Various studies have found that some pathogens are seasonal, peaking at different times of the year [4–6]. Frequently, the seasonal periodicity of diarrheal disease is attributed to weather, which could drive incidence by diverse mechanisms. For example, weather conditions can favor the survival and replication of pathogens on fomites [7], the transmission between human hosts through flooding and contamination of drinking water [8], and the prevalence of vectors that transmit disease between hosts [9,10]. Weather has broadly been shown to be mathematically correlated with diarrhea incidence [11,12], with some computational studies claiming a causal link [13] despite potential limitations to their methodology [14–16].

However, most studies of disease seasonality have been conducted in temperate climates, and substantially less is known about the seasonality of diseases in tropical countries [17,18], where diarrheal disease is one of the leading causes of morbidity and mortality among children [19]. The wide variety of climates and populations in the tropics make it challenging to uncover general patterns in the epidemiology of diarrheal disease. Compounding these challenges, most studies are limited to sites within a single country focused on a specific disease. Characterizing the seasonal epidemiology of these pathogens could enable clinicians to better diagnose children based on the time of the year. Additionally, identifying the weather conditions associated with each pathogen could help us infer pathogen transmission pathways, predict large outbreaks, and develop intervention strategies.

In this article, we perform a secondary analysis of the Global Enteric Multicenter Study (GEMS), a large, multi-country study of moderate-to-severe diarrhea (MSD) among children younger than five years of age [2], to investigate the underlying patterns of pathogen-specific seasonality for diarrheal disease in resource-limited settings. We focus on the pathogens associated with the plurality of attributable diarrheal illnesses in GEMS: rotavirus, *Cryptosporidium, Shigella,* typical EPEC (tEPEC) and enterotoxigenic *Escherichia coli* encoding heat-stable enterotoxin (ST-ETEC), adenovirus 40/41, and *Campylobacter* [2,20]. Utilizing a variety of mathematical techniques, we analyze the GEMS data to measure the strength of pathogen seasonality at each site, to link local weather conditions to pathogen prevalence, and to reveal site-specific seasons associated with pathogens. The data from GEMS affords a rare opportunity to compare the seasonality of many pathogens across sites in different countries using data from a single coordinated study.

## Materials and methods

### GEMS data

The Global Enteric Multicenter Study (GEMS) was conducted for 36 consecutive months from December 2007 to March 2011 at sites in seven communities in sub-Saharan Africa (Basse, The Gambia; Bamako, Mali; Nyaza Province, Kenya; Manhiça, Mozambique) and south Asia (Karachi [Bin Qasim Town], Pakistan; Kolkata, India; Mirzapur, Bangladesh) [2]. Children 0–59 months of age with moderate-to-severe diarrhea (MSD) who lived inside of the site’s enumerated catchment area were eligible to enroll. For a child to be included, the diarrhea episode needed to be new (onset after at least seven days without diarrhea), acute (onset within seven days), and had to meet at least one clinical criteria for MSD. Clinical criteria for MSD included the following: the child presents clinical signs of dehydration, i.e., sunken eyes or loss of skin turgor, assessed by clinician and confirmed by mother, prescription or use of intravenous hydration; dysentery identified by blood in stool; or admission to the hospital for diarrhea or dysentery [2]. To limit the number of enrollments and ensure balanced enrollment by age, 8–9 children in each age strata (0–11 months, 12–23 months, 24–59 months) were recruited each fortnight (14 days) at each site. Stool samples from enrolled children were tested for pathogens. In total, GEMS enrolled 9,439 out of 14,753 eligible children with MSD during the study period (Table 1). Matched controls, who did not have diarrhea, were not included in our analysis.

**Table 1.**
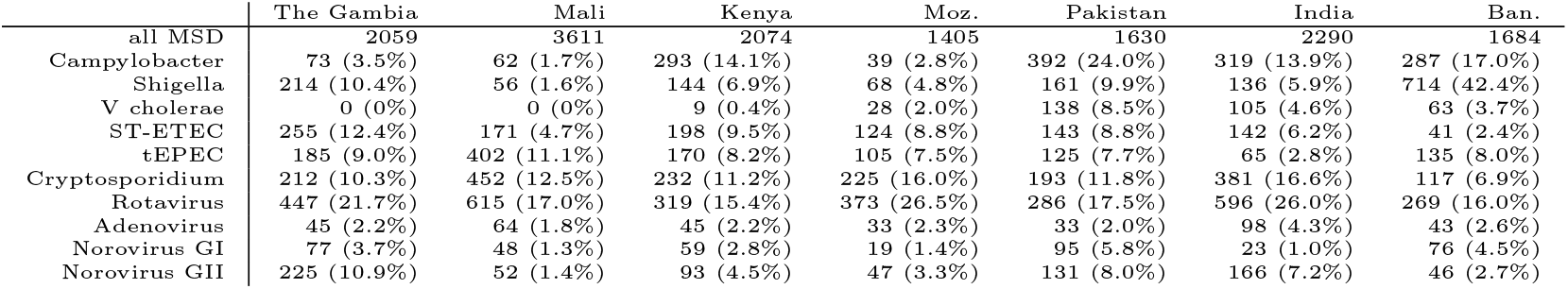
Estimated prevalence of pathogens among children with MSD. We estimated the number and percentage of eligible children positive for each pathogen based on the proportion of tested cases who were positive.

We limited our analyses to pathogens associated with the plurality of attributable diarrheal illnesses in GEMS [2]: rotavirus, Cryptosporidium, *Shigella* spp., typical EPEC (tEPEC), and enterotoxigenic *Escherichia coli* encoding heat-stable enterotoxin with or without genes encoding heat-labile enterotoxin (ST-ETEC). Two additional major pathogens identified in a subsequent analysis were included [22]: adenovirus 40/41 and *Campylobacter* spp. We also included three pathogens that were identified as prevalent in multiple sites during at least one of the seasons: *V. cholerae,* norovirus GI, and norovirus GII.

### Estimating pathogen prevalence among all eligible visits for two week periods

To estimate the number of children who would have tested positive for a pathogen had all eligible children been enrolled, we assume that the proportion of children testing positive for a pathogen is equal among enrollees and those eligible within age strata (0–11, 12–23, and 24–59 months old) and fortnight of clinic visit. Fortnights are defined as consecutive 14-day periods starting with the first enrollment at each site. For example, Fig. 1(A) illustrates the difference between rotavirus confirmed enrolled children (purple) and the total estimated rotavirus prevalence among eligible children (black lines) in Bangladesh. Thus, the overall estimated pathogen prevalence includes the laboratory-positive children among the enrolled plus the estimated proportion among the eligible, reflecting the true age distribution of MSD cases. Fig. 1(A) does not show a large difference between confirmed rotavirus cases and estimated rotavirus cases due to the small differences between the number of eligible and enrolled children. Fig. S1 for the Mali study site illustrates a wider difference between the number of eligible and enrolled children, resulting in a wider discrepancy between confirmed and estimated rotavirus cases.

**Fig 1.**
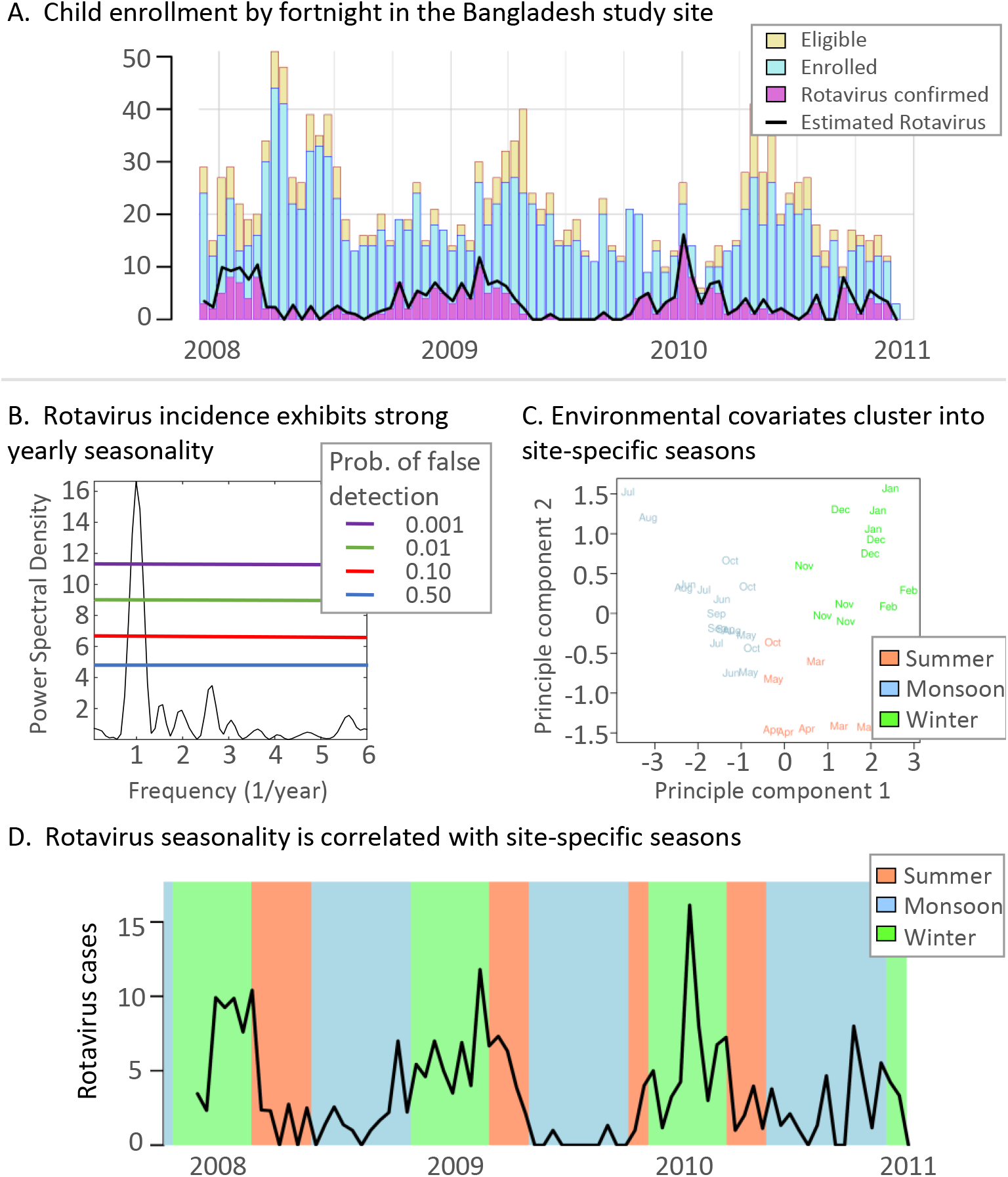
Analysis overview. A. The number of eligible and enrolled children for the Bangladesh study site each fortnight (blue and gold, respectively). The number of rotavirus-confirmed samples from enrolled children are in purple. The estimated rotavirus prevalence among all eligible children which includes confirmed positive and estimated positive from the eligible children is illustrated by the black line. B. The power spectral density of the estimated rotavirus positive prevalence time-series. Each colored line indicates the probability of a random time-series signal with the same length and sampling being misinterpreted as a true signal. C. Seasons were identified using PCA to reduce the dimensionality of the weather variables and k-means clustering. The color of each month corresponds to the cluster and season. For Bangladesh, we call each season by the colloquial names summer, monsoon, and winter. D. The estimated number of cases positive for a pathogen each fortnight (black line). Each background color corresponds to the data-driven season identified in C.

### Weather data for GEMS study sites

Daily historical weather data from weather stations were acquired from the Global Surface Summary of the Day (GSOD) [21]. These data include daily measurements of temperature, precipitation, and dew point. For some GEMS study sites, the closest weather station does not have complete data for the entire study interval (2007–2011). In these cases, we choose the closest weather station with nearly complete data; for example, the weather station for The Gambia site is in Tambacounda, Senegal which is approximately 77 kilometers away and is the largest distance between site and station used in this study. Both the raw data downloaded from [21] and our filtered data files for the analysis can be found at [22].

Relative humidity was computed from the GSOD using the following empirical relationship, originally defined by Bosen in 1958 [23]:

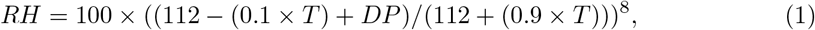

where T is temperature in Celsius, *DP* is dewpoint in Celsius, and RH is relative humidity in percentage. We estimated specific humidity:

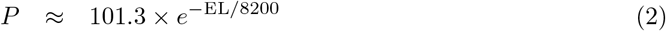

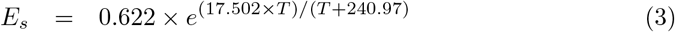

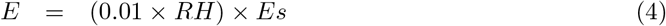

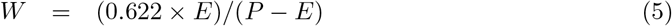

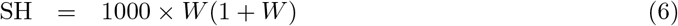

where EL is elevation in meters, *P* is air pressure in kPa, *E_s_* is saturation vapor pressure in kPA, *E* is the vapor pressure in kPa, *W* is the mixing ratio, and SH is the specific humidity in g/kg [personal communication from James Tamerius]. Fig. S4 shows the temperature, rain, and specific humidity for each GEMS site during the study period. Days with missing rainfall data were assumed to have no rainfall, and days with missing temperature or humidity data were not included when computing average temperature or humidity for the fortnight or month containing them.

### Detecting periodicity and relative phase in disease incidence time-series

Seasonality was identified by estimating the power spectral density contained in the estimated incidence time-series data for each pathogen separately. Specifically, we utilize the Lomb-Scargle periodogram [24–26] to identify the spectral power for each resolvable frequency, which is constrained by the temporal length of GEMS and our two-week sampling interval. The Lomb-Scargle periodogram is especially useful for analyzing constrained time-series; the technique quantifies the uncertainty of detecting a specific periodicity [27] even with unevenly sampled data [25]. Fig. 1(B) shows the standard Lomb-Scargle periodogram which is scaled by two times the variance of the pathogen time-series. Each horizontal line indicates a power-level threshold providing the probability of a random signal being mischaracterized as a true periodic signal. We quantify the uncertainty of the periodicity of a time-series as a probability of false detection denoted as *p_fd_* < threshold. We used the MATLAB function *plomb* from version 2014A to estimate the spectral density; see [22] for all computational scripts.

For each combination of pathogen and site, we used the fast Fourier transform (FFT) to extract the relative phase of when each pathogen annually peaks. The FFT and periodogram are mathematically connected. Here, we use the FFT to identify the relative timing of the peak of disease incidence across diseases. We performed this transformation with respect to the first day in the two-week period. Computing the phase from the FFT of the disease incidence time-series was implemented in MATLAB version 2014A; the computational scripts can be found at [22].

### Identifying associations between environmental factors and pathogens

We estimated the strength of association between weather covariates (i.e., cumulative precipitation, average temperature, average relative humidity, and average specific humidity) over the fortnight and the estimated number of children positive by pathogen and by country. Specifically, we identify the highest and lowest quartiles for the number of children with MSD estimated to be positive by fortnight. Quartiles were chosen due to low sample numbers for certain pathogens; for example, some pathogens were detected in only 1/4 of the fortnights. In these cases, the median could not be used to distinguish high and low prevalence, and the weather during fortnights with no positive cases was compared to the weather during fortnights with cases. The associations between pathogen prevalence and environmental values are computed using a Wilcoxon rank-sum test. The computations were performed in the R scientific computing environment and the code can be found in [22].

### Identifying data-driven seasons directly from weather data

We identify seasons directly from the monthly weather data derived from GSOD, instead of using pre-defined seasons from the scientific literature. These seasons are *data-driven* in the sense that our algorithm clusters months that have similar weather covariates. We transform daily environmental values to monthly values by summing rainfall and taking the average values for temperature, relative humidity, and specific humidity. However, the yearly variability has a substantial impact on the beginning and end of seasons; see Fig. S4 for an illustration of the season variability when using actual weather data to identify seasons.

To determine seasons from weather data, we used principal component analysis (PCA) to reduce the number of weather covariates from four to three. The dimensionality reduction with principal components analysis eliminates redundant variables [28]. We implemented k-means clustering to group weather-months by similarity; three clusters best described the groupings of weather-months. Fig. 1(C) shows each of the month’s weather data projected on the leading two principal components colored by cluster. We call these clusters data-driven seasons. This procedure identified reasonable quantitative definitions of seasons that qualitatively matched descriptions for each site and are broadly consistent with the literature. Some seasons can occur more than once per year, e.g., there could be two rainy seasons in one calendar year. Fig. 1(D) illustrates the timing of seasons in Bangladesh. We used the R scientific computing environment and provide the analysis code at the following repository [22]

### Determining significant differences in pathogen-positive cases by season

To determine if the estimated number of children positive for pathogens each month had significant differences among our data-driven seasons, we used the Kruskal-Wallis test. We then used Dunn’s test to identify the pairs of seasons with significantly different numbers of children positive for pathogens. Both statistical tests help characterize the association of MSD with weather and seasons. The tests were applied using the R scientific computing environment.

### Ethics approval

The GEMS study protocol was approved by ethics committees at the University of Maryland, Baltimore and at each field site. Parents or caregivers of participants provided written informed consent, and a witnessed consent was obtained for illiterate parents or caretakers.

## Results

### Annual and biannual cycles are detected for several diarrheal pathogens and study sites

Fig. 2 plots the number of study-eligible children (children with MSD) visiting the clinics each fortnight and the estimated number of them positive for rotavirus, *Cryptosporidium,* and *Shigella.* The number of children with MSD in Mali, Mozambique, Pakistan, India, and Bangladesh had significant annual periodicity *(p_fd_* <10%) (Fig. 3). We studied the periodicity of the estimated number of eligible children positive for eight pathogens: *Campylobacter, Shigella,* ST-ETEC, tEPEC, *Cryptosporidium,* rotavirus, and adenovirus 40/41. *V. cholerae,* norovirus GI, and GII did not have statistically significant annual seasonality at any of the sites. Each of the seven sites had 1-5 pathogens with significant annual periodicity *(p_fd_* <10%). Rotavirus has significant annual periodicity in The Gambia, Mali, India, and Bangladesh (all *p_fd_* <1%). *Cryptosporidium* has significant annual periodicity *(pf_d_* <1%) in The Gambia, Mali, and Mozambique. *Shigella (p_fd_* <0.1%) and EAEC (*p_fd_* <1%) had significant annual periodicity only in Bangladesh. ST-ETEC and tEPEC had significant annual periodicity at *p_fd_* <1% only in Mozambique. We also tested for 6-month periodicity (5.5-6.5 month period) using the same procedure as above. Out of the seven sites, only Kenya had pathogens with a significant 6-month periodicity: rotavirus and norovirus GII *(p_fd_* <10%).

**Fig 2.**
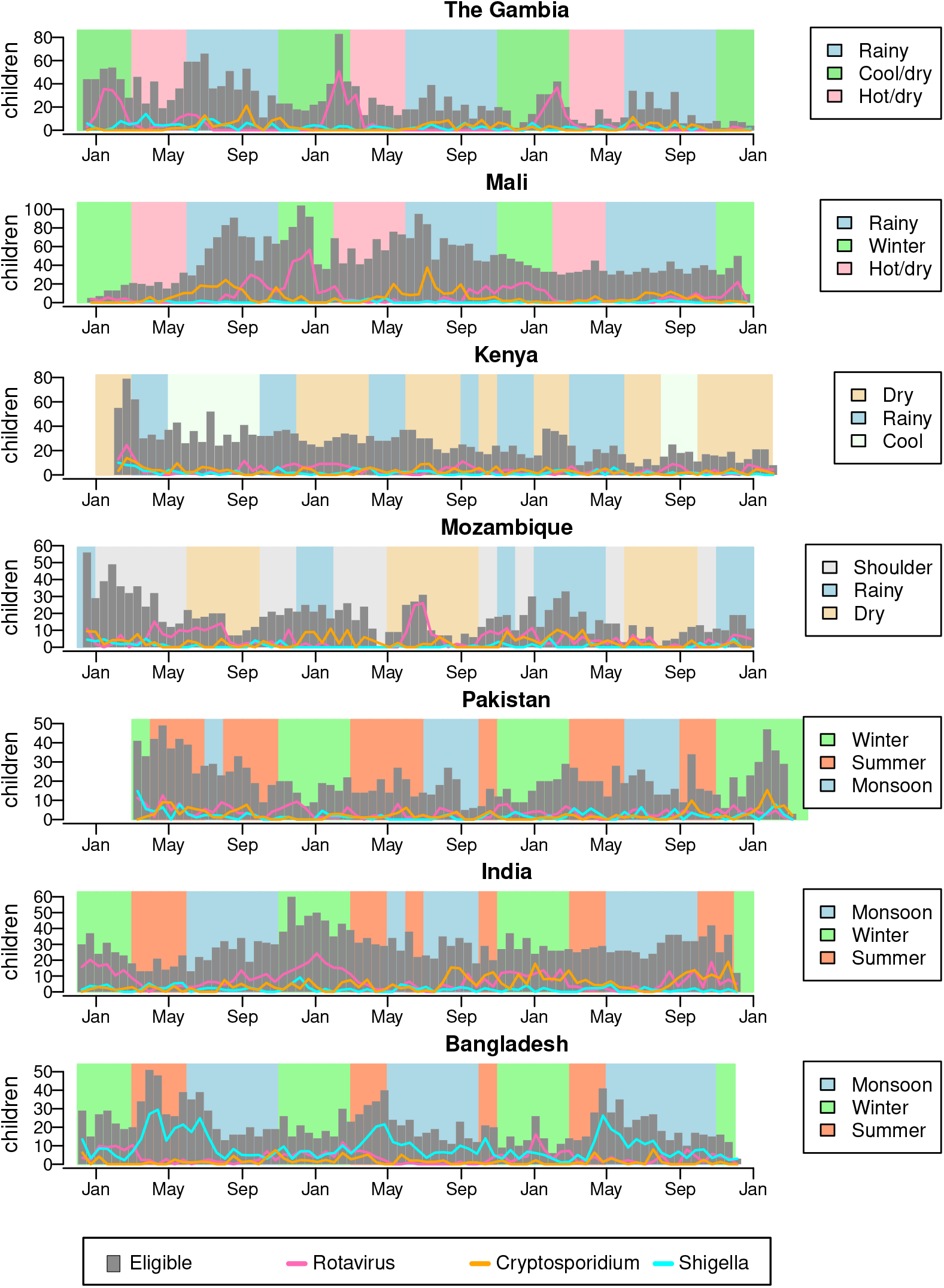
Estimated fortnightly cases positive for rotavirus, *Shigella,* and *Cryptosporidium.* Gray bars plot the number of children with MSD visiting the study clinics each fortnight. The lines show the estimated number of these children positive for rotavirus, *Shigella,* and *Cryptosporidium.* The background shading indicates the timing of data-driven seasons at each site.

**Fig 3.**
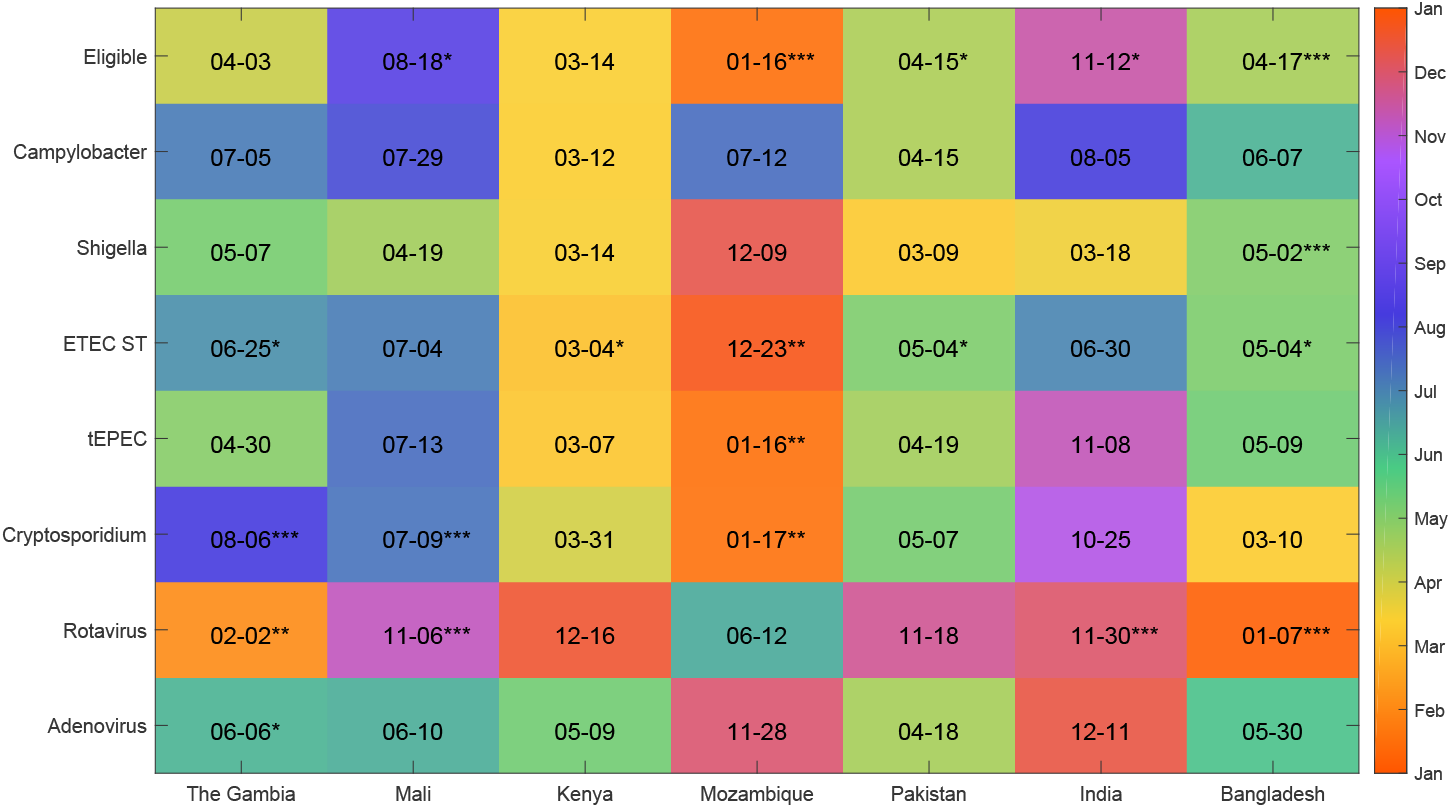
Annual peak dates of pathogens. We used the phase information from the FFT of the number of eligible children estimated to be positive each fortnight to determine the date of the annual peak of each pathogen in each country (indicated as month-day in each square and color-coded by day of year). The significance (false-detection rate *p*¡_d_) of the annual cycle was obtained from the Lomb-Scargle periodogram of the most significant period between 11 and 13 months, indicated by **** for <0.1%, *** for <1%, ** for <5%, and * for <10%.

### Broad statistical association between fortnightly pathogen prevalence and weather

We found that fortnights with a high number of MSD clinic visits tended to have significantly higher rainfall and humidity in Mali and Mozambique than fortnights with a low number of MSD visits (Wilcoxon rank-sum test *p* <0.05). In contrast, fortnights with a high number of MSD visits were less rainy in India and less humid in The Gambia. Fig. 4 illustrates the statistically significant associations between weather and pathogen prevalence by GEMS site. There were substantially more statistically significant associations between weather covariates and pathogen prevalence than detection of annual periodic cycles; see Figs. 3 and 4 for a visual comparison across site and pathogen. We also detected a significant association between increased numbers of estimated rotavirus positive cases and lower humidity, rain, and temperatures. Hot weather was also associated with high numbers of *Shigella*-positive cases (Fig. 4). ST-ETEC, tEPEC, and *V. cholerae* were associated with hot and humid weather. Rainy and humid fortnights had significantly more *Cryptosporidium-positive* cases than dry fortnights.

**Fig 4.**
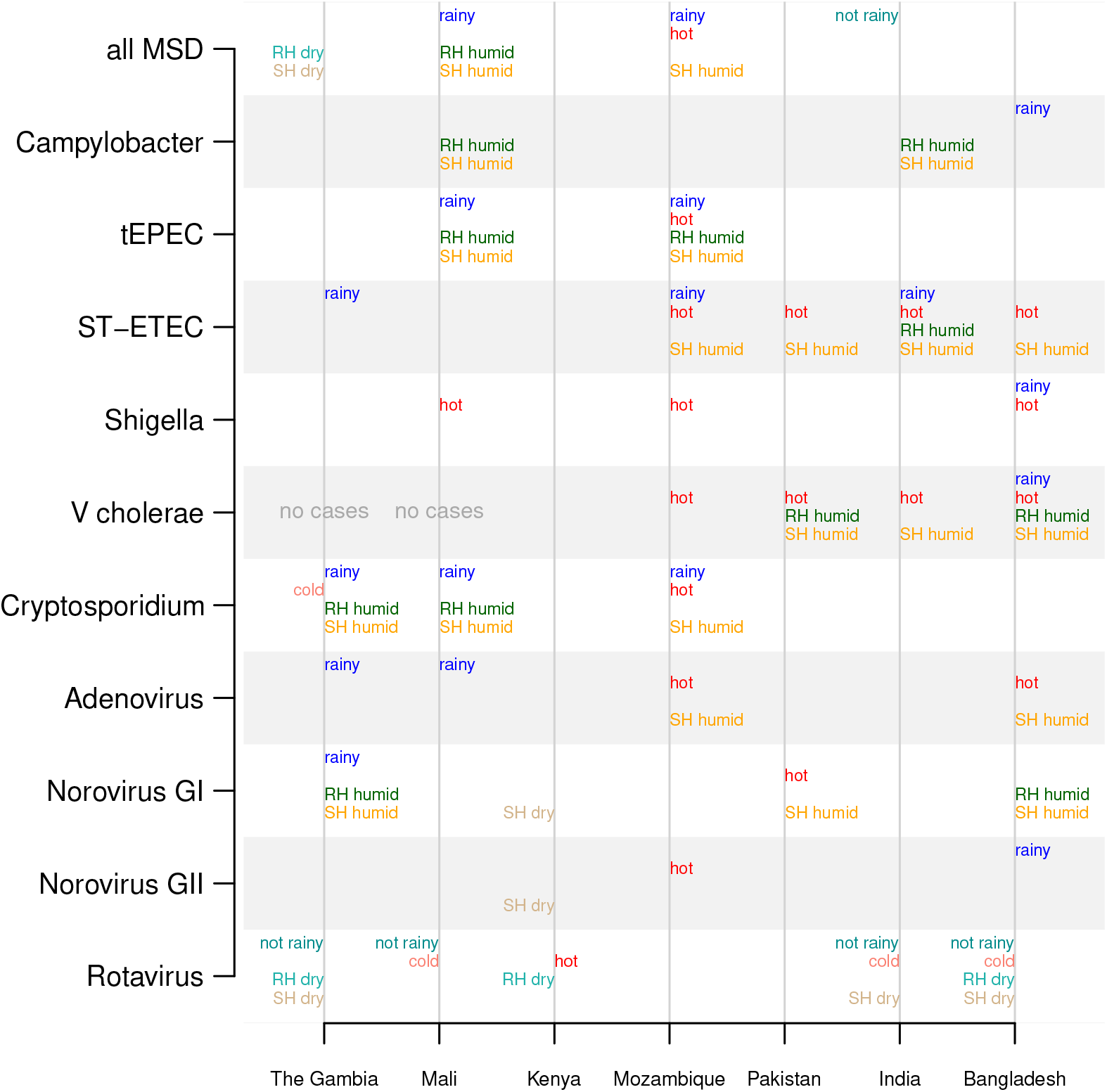
Significant differences in weather of fortnights with high vs low pathogen prevalence. The weather covariates with significant differences (p<0.05 using the Wilcoxon rank-sum test) between fortnights with high vs low prevalence of pathogens are printed in the chart. Weather covariates that are higher during high-prevalence weeks are printed to the right of the vertical lines (“rainy” indicates higher rain in pathogen-associated fortnights, “hot” means higher temperatures, “RH humid” means higher relative humidity, and “SH humid” means higher specific humidity), and those that are lower during high-prevalence weeks are printed on the left (i.e., “not rainy”, “cold”, “RH dry”, and “SH dry”).

### Site-specific seasons can be identified from weather data

We identified three data-driven seasons for each GEMS site. Fig. 1(C) illustrates the clustering of monthly weather covariate data, collected from the closest weather station in Bangladesh during 2008 to 2011, projected on to the first two principal components.

The color of each month indicates the cluster *(season)* to which the month belongs. Table S1 summarizes average weather conditions for each season at each site. Fig. 1(D) illustrates how the clustering is reflected temporally during the study time period. These data-driven seasons broadly fit informal definitions of seasons within each study site. This methodology, however, allows for seasons to vary in duration and initiation time and is based on site-specific weather data from the duration of study enrollment (Fig. S4). Even when sites share the same names for season (e.g., summer and winter), the actual weather conditions for these seasons differ but the relationships among seasons is consistent (e.g., summer is hotter than winter, and rainy season has more rain than dry season) (Table S1).

### Pathogen prevalence is strongly associated with data-driven seasons

We found rotavirus tended to have significantly higher prevalence in winter or cool/dry seasons than during rainy seasons (Fig. 5). For example, the estimated rotavirus prevalence in The Gambia was statistically different between the rainy and two other seasons, but not statistically different between the cool/dry and hot/dry season. We did not find a country-pathogen combination with statistical different prevalence among all three seasons. We found *Shigella* was most prevalent during the summer in Bangladesh and the hot/dry season in Mali (Fig. 5). ST-ETEC was most prevalent during the rainy season in The Gambia and Mozambique and the summer and monsoon seasons in Bangladesh and India. *Cryptosporidium* had highest prevalence in the rainy season in The Gambia, Mali, and Mozambique. Bacterial pathogens and *Cryptosporidium* had lower prevalence in winter or cool/dry seasons than in rainy or summer seasons.

**Fig 5.**
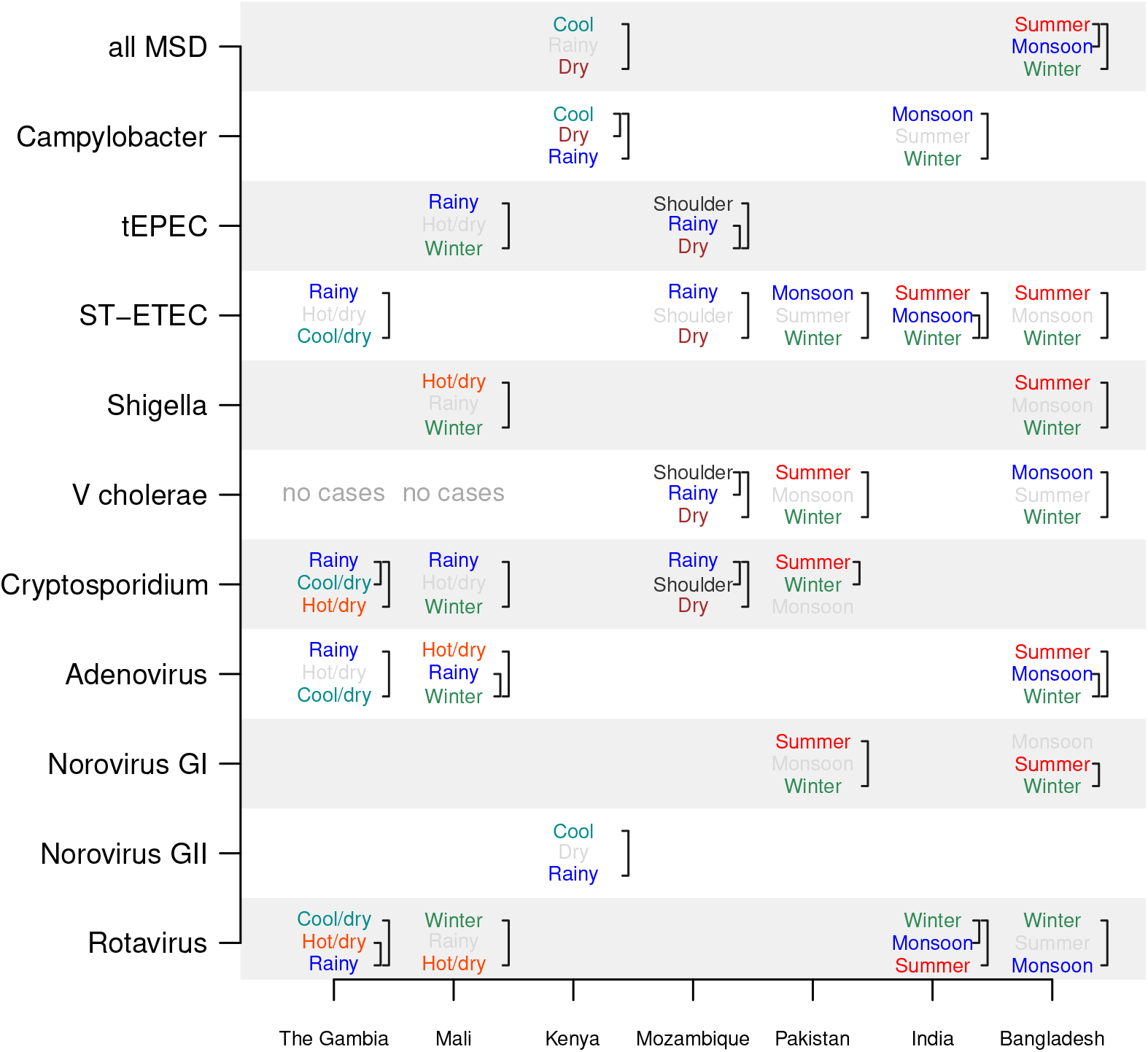
Relative prevalence of pathogens across seasons. The Dunn test is used to compare the monthly cases between pairs of seasons. If there is a significant difference between seasons, their names are printed, with the season with the most cases per month on top and fewest on the bottom. Brackets indicate pairs of seasons with significant differences, and season names are color-coded (summer, winter, monsoon, hot/dry, cool/dry, etc) when there is a statistically significant difference between seasons (p<0.05 using a Bonferroni correction) and light gray otherwise. No country–pathogen combination had significant prevalence differences among all three seasons.

### The most prevalent pathogens differ by season and child age

Because different pathogens were associated with different seasons, the relative prevalence (and ranking) of pathogens among children with MSD differed by season. Among children 0-23 months old, rotavirus was the pathogen most frequently detected among eligible children in the winter, dry, or cool/dry seasons at all sites (22-57% of cases, Fig. 6, left panel). In the rainy or monsoon months, rotavirus was less dominant, and *Shigella, Campylobacter,* and *Cryptosporidium* were the most often detected. Among those 24-59 months old, rotavirus was the most frequently detected pathogen at three of the seven sites in winter, dry, or cool/dry seasons (Fig. 6, right panel). For these older children, *V. cholerae* is one of the top three pathogens at some sites during the monsoon or summer seasons.

**Fig 6.**
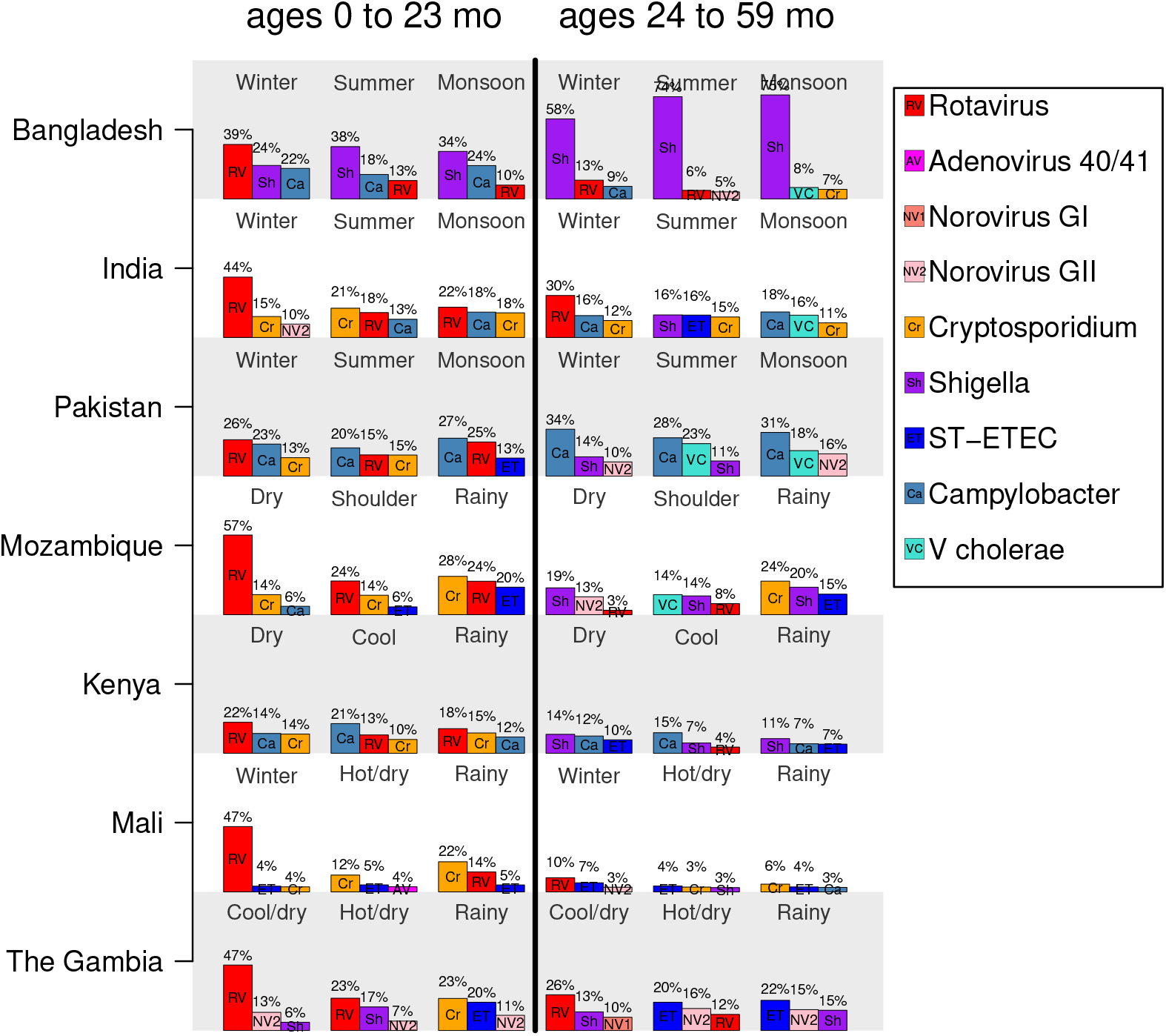
The top three pathogens by season. For each country, the top three most frequently detected pathogens (out of the nine considered) among eligible children ages 0 to 23 months (left half of the plot) and 24 to 59 months old (right half) are shown. Bar heights are proportional to the percent of cases positive for a pathogen (which is also printed above each bar), and the pathogens are indicated by abbreviations and color as summarized in the legend. Pathogens considered are: rotavirus, adenovirus 40/41, norovirus GI, norovirus GII, *Cryptosporidium, Shigella,* ST-ETEC, *Campylobacter,* and *V. cholerae.* Note that the seasons are not plotted in a particular order and may be non-sequential.

### Annual cycles indicate a birth month is a risk factor for diarrheal illness

If MSD or pathogen associated with MSD has a highly significant annual period, the age at which a child is most likely to get diarrhea depends on the birth month of a child. For example, if diarrheal disease peaks in the first quarter of the year, a child born in the second quarter might not be at high risk until six to nine months of age, while a child born before the first quarter could be at high risk of exposure soon after birth. This effect can be seen in The Gambia, as shown in the heat maps of birth month vs age of MSD visit shown in Fig. S2. Strong seasonality is visible as diagonal “stripes” in these heatmaps. Similar effects can be seen among rotavirus-positive MSD visits when rotavirus has a significant annual periodicity (Fig. S3).

## Discussion

In a systematic manner, we have characterized the seasonality of diarrheal pathogen prevalence at seven study sites in the Global Enteric Multicenter Study (GEMS). The GEMS study provided an unprecedented perspective on the population-based burden of moderate-to-severe diarrhea among children in resource-constrained settings spanning multiple countries and continents [2]. In this work, we leverage study data collected over a three year period to better understand the seasonality of the most important diarrhea-associated pathogens as well as the relationship of prevalence to site-specific environmental weather variables. We utilize standard mathematical methodologies to test for seasonality as well as more modern machine-learning algorithms to identify data-driven seasons from detailed environmental data to correlate with the periodicity of pathogen incidence. This work provides a unique perspective on characterizing the seasonal epidemiology of diarrheal pathogens due to the diversity of study sites in resource constrained settings, consistent study protocol, and large sample size.

Rotavirus was the only pathogen with highly significant annual prevalence peaks at most of the seven study sites. Previous studies have found that rotavirus diarrhea is seasonal and tends to occur in the winter, but these study sites have been primarily in temperate regions [4–6]. Recent studies also suggest rotavirus seasonality might be weaker in the tropics, with some evidence that it peaks during the cooler seasons [4–6]. We found less consistent periodicity of other pathogens across GEMS sites; however, there is less evidence in the scientific literature on the global seasonality of diarrheal pathogens other than rotavirus. We hypothesized that weather drives the transmission of many diarrheal pathogens at GEMS sites, and the lack of annual periodicity in different GEMS sites or by pathogen may be the result of a milder and more variable weather drivers such as those found in tropical regions.

Despite the lack of evidence of annual periodicity of pathogen prevalence time-series, we found that diarrheal pathogen prevalence is associated with weather. Several bacterial pathogens were more prevalent during hot and rainy weather, which could favor the growth of bacteria in the environment or the contamination of water sources [29]. We found ST-ETEC was generally associated with warmer weather (in Mozambique, Pakistan, India, and Bangladesh), consistent with the previously observed association between ETEC and higher temperatures and not rainfall [30, 31]. A significant association between *Cryptosporidium* and rainy weather was identified in The Gambia, Mali, and Mozambique. In contrast, rotavirus was found to be more prevalent during the drier winter months out of phase with *Cryptosporidium*; this is broadly consistent with reviews of rotavirus seasonality studies in the tropics [5,32]. Note, however, that other studies have found peaks of rotavirus activity during monsoon seasons in some tropical sites [33,34]. Our results for Bangladesh generally agree with the analysis by Das et.al. [35] where they found rotavirus peaked in the winter, cholera in the monsoon, and ETEC in the summer at three sites in Bangladesh. However, it is difficult to identify the combination of components of weather that drives each pathogen, since weather covariates can be highly correlated (e.g., heat and humidity).

We found that the weather conditions at each site seemed to fall into a few distinct classes, such as “warm, humid, and rainy”, “hot and dry”, and “cool and rainy”, and these classes broadly agree with informally identified site-specific seasons, such as “summer” and “winter”. We used modern machine-learning techniques to formalize this observation and cluster the environmental data by site to identify data-driven seasons. To our knowledge, associating pathogen prevalence with data-driven seasons is an innovation for revealing site-specific weather trends that are potential drivers of pathogen prevalence. We found that pathogen prevalence was often significantly different across seasons, and that pathogens might lack strict annual periodicity because of the year-to-year variability in the timing and length of seasons. We believe that these seasons serve several purposes when studying pathogen association with weather conditions: 1) Seasons last for a few months while weather conditions can change daily so associations with seasons are more robust to disease reporting lags and frequency; 2) Some pathogens may be driven by a combination of weather conditions (e.g., heat and humidity), which are captured by seasons but not by individual weather covariates; and 3) Seasons give us a common terminology to use across different sites.

These site-specific seasons enable public health officials and clinicians to parse out population pathogen prevalence changes across seasons; Fig. 6 shows how the top three pathogens change across seasons, sites, and age of child. Among children with MSD 0–23 months of age, rotavirus was the pathogen most frequently detected, particularly during winter or cool/dry seasons. *Cryptosporidium* was the top pathogen among 0-23 month-olds during the rainy season in some countries. Among children 24–59 months old, *Campylobacter, Shigella,* and ST-ETEC were the most frequently detected pathogens. This detailed site-specific description could help with differential diagnosis and treatment choices with diarrheal symptoms if laboratory services are unavailable.

For sites with strong annual rotavirus seasonality, the birth month of a child is associated with age-dependent risk of rotavirus diarrhea. A similar result was reported in a study of rotavirus in England and Wales, a high-income country, where children born in the summer had a higher risk of rotavirus diarrhea in the first year of life compared to those born in the winter [36]. Broadly, this data and analysis can used to assess risk of a diarrheal disease by birth month; from a public health perspective, these results could be included in supply chain and operational planning for clinics and hospitals.

The smaller seasonal changes in temperature and humidity in some tropical settings compared to temperate ones could make it difficult to study weather as a driver of diarrheal disease. We found the association between pathogens and weather is less pronounced at sites with less seasonal variation in weather, such as the study sites in Kenya and Mozambique. Because temperate and tropical climates have different ranges of weather conditions, the relationship between pathogens and weather covariates could differ [37]. Shorter seasons may make detecting the association between weather and pathogens difficult, since the pathogen would have less time to respond (e.g., amplify, transmit) to changes in weather conditions. The Kenya site has two rainy and two dry seasons per year, thus weather covariates had bi-annual rather than annual periodicity, and rotavirus and norovirus GII prevalence had significant bi-annual periodicity, but previous studies noted that the seasonality of rotavirus disease is subtle in Kenya [38]. Weather might also mediate complex transmission pathways [12]; for example, *Shigella* had been observed to peak in April–June at the GEMS study site in Bangladesh and was associated with seasonal peaks in housefly density in February and March [39]. Therefore, peaks in disease prevalence could be driven by the weather during a preceding season. Weather’s effects on pathogen transmission may also interact with population density, so that adjacent urban and rural areas can experience differing pathogen seasonality [33], which could make it difficult to generalize the relationships between weather and pathogen prevalences.

It is mathematically challenging to determine if a pathogen is causally associated with seasonal environmental factors. This study used three years of observations, so variation in weather between years could mitigate the confounding of climate and calendar-based covariates [11,12]. We determined associations between weather and pathogen prevalence providing statistical support for our conclusions; moreover, we describe the probability of mis-identifying a seasonal signal by site and pathogen, based on the study duration and surveillance sampling. We did not, however, attempt to identify *causally* linked environmental factors. There is the potential that additional years of data or spatial weather covariates within sites could further improve our ability to link weather to disease prevalence. More sensitive pathogen detection assays could also improve our ability to analyze seasonality. A recent reanalysis of a subset of the samples from GEMS revealed higher prevalence of some pathogens among cases, particularly *Shigella,* adenovirus 40/41, ST-ETEC, and *Campylobacter* [20]. Even with more sensitive assays and longer time series, determining the etiology of diarrheal disease is difficult, since multi-pathogen infections are common and disease could be caused by (or even mitigated by) one of the pathogens. We primarily focused on pathogens that have strong associations with diarrhea (e.g., low minimum infectious dose) to mitigate this challenge. One result of this work is to provide caution to the global health community, especially given the current trend of estimating burden at a fine-scale spatial resolution with an underlying statistical model that relies too heavily on data from a nonrepresentative region.

Notwithstanding many of these limitations, our study was unique in studying the seasonality of multiple pathogens across multiple countries at the same time using the same study design. Although the primary purpose of GEMS was to identify the most prevalent and virulent pathogens associated with MSD at the study sites, the association of certain pathogens with weather covariates was strong enough to study. We believe that identifying the environmental conditions that facilitate transmission of pathogens could help us understand the mechanisms by which they spread in human populations and choose the most effective interventions to reduce transmission [40–43]. We believe that this study will better inform the global health community around pathogen prevalence in different resource constrained settings. Moreover, the identification of age-dependent risk of pathogen and population prevalences by season could lead to better clinical diagnoses and allocation of public resources.

## Supporting Information Legends

**S1 Checklist: STROBE checklist of items for case—control studies**

**Table S1. Average monthly conditions of regional seasons.** Months indicate when these seasons have occurred during the study, which can vary slightly by year. “% time” is the % of the study months assigned to each season.

**Fig S1. Estimating prevalence of rotavirus among children with MSD.**

Children with MSD were eligible for the study (gold bars). A sample of them were enrolled in GEMS each fortnight (blue bars) and stools samples were tested for rotavirus and other pathogens. The estimated number of eligible children positive for a rotavirus (black line), is extrapolated from the positivity among enrolled children (purple bars) during the same fortnight.

**Fig S2. Age and birth month distribution of eligible children.** The numbers in the plot are the number of eligible children born in month x and visited the clinic at age y. Yellow areas are low numbers of eligible children, while red numbers are high. Note the diagonal elements in countries where there is a distinct diarrheal disease season.

**Fig S3. Age and birth month distribution of enrolled cases who were rotavirus-positive.** The numbers in the plot are the number of enrolled cases born in month x and were enrolled at age y who were rotavirus-positive. Note the diagonal elements in countries where there is a distinct rotavirus season. Enrollment rates could differ by age group, so one should not compare numbers across age groups (0-11m, 12-23m, 24m and older).

**Fig S4. Seasons and monthly temperature, rainfall, and specific humidity for each site from 2007—2011.** Seasons were defined by a k-means clustering algorithm on PCA-transformed monthly weather covariates. Relative humidity not shown.

Author summary
Diarrheal disease is one of the leading causes of death among young children in the developing world. It is difficult to determine which of a wide variety of pathogens are most responsible for disease, since this differs by location and time of year. Here, we study the seasonal prevalence of several pathogens among children with moderate-to-severe diarrhea across study sites in Africa and South Asia. We found that several pathogens, including rotavirus, had regular annual peaks. Some pathogens were associated with weather conditions, such as heat or rain, or with general seasons of the year, such as summer or winter. We believe that describing the seasonal epidemiology of these pathogens could enable better diagnoses of symptomatic children based on the time of the year. Additionally, weather is a major driver of diarrheal pathogen transmission, and identifying the conditions associated with each pathogen could help us infer pathogen transmission pathways, predict large outbreaks, and develop intervention strategies.

